# A Y chromosome-linked genome editor for efficient population suppression in the malaria vector *Anopheles gambiae*

**DOI:** 10.1101/2024.05.14.594116

**Authors:** Ignacio Tolosana, Katie Willis, Austin Burt, Matthew Gribble, Tony Nolan, Andrea Crisanti, Federica Bernardini

**Affiliations:** Department of Life Sciences, Imperial College London, London, UK; Department of Vector Biology, Liverpool School of Tropical Medicine, Liverpool, UK

## Abstract

Genetic control – the deliberate introduction of genetic traits to control a pest or vector population – offers a powerful tool to augment conventional mosquito control tools that have been successful in reducing malaria burden but that are compromised by a range of operational challenges. Self-sustaining genetic control strategies have shown great potential in laboratory settings but hesitancy due to their invasive and persistent nature may delay their implementation. Here instead we describe a self-limiting strategy, designed to have geographically and/or temporally restricted effect, based on a Y chromosome-linked genome editor (YLE). The YLE comprises a CRISPR-Cas9 construct that is always inherited by males yet generates an autosomal dominant mutation that is transmitted to over 90% of the offspring and results in female-specific sterility. Males are unaffected. To our knowledge, our system represents the first engineering of the Y chromosome to generate a genetic control strain for mosquitoes. Mathematical modelling shows that this YLE technology is up to 8 times more efficient for population suppression than optimal versions of other self-limiting strategies.

## INTRODUCTION

Vector control has proven to be one of the most efficient ways to reduce malaria incidence through history^1–3^. The use of insecticide treated bed nets (ITNs) and indoor residual spraying (IRS) have driven the reduction of malaria cases since the year 2000^1,2^. However, different factors such as challenges in the use and distribution of bed nets, the rise of resistance towards insecticides and antimalarial drugs, and the behavioural plasticity of the mosquito vectors have contributed to the decrease in the progress against malaria since 2015, highlighting the need for new tools and strategies to control the disease^2^.

Genetic control of vector populations arises as a very promising field in this framework, as it offers a range of technologies with diverse efficacy and control prospects, but with the common characteristic of being species-specific and environmentally non-polluting^4,5^. These strategies can be divided into those that aim at modifying the mosquitoes’ genome to prevent the development of the malaria parasite^6,7^ and those that aim at suppressing mosquito populations^8–11^. Genetic approaches for vector control also differ in their intended post-release dynamics: some technologies are designed to increase in frequency from a small release and spread geographically (self-sustaining strategies), while others pose a temporal and spatial restriction on the persistence of the modification (self-limiting) and require large and continuous releases to maintain their effect on the target population^4,12^.

Traditionally, the Sterile Insect Technique (SIT) has relied on the release of radiation-sterilised males, and it has been widely applied for insect pest control programmes^13–15^. More recently, transgene-based SITs and other alternative self-limiting technologies based on the presence of lethality-inducing transgenes such as the Release of Insects carrying a Dominant Lethal gene (RIDL), and on synthetic sex ratio distortion systems capable of inducing a strong male bias in the progeny have been developed and tested in the lab or in the field for different vector species^10,16–21^. While these strategies have been useful in certain contexts, they share the crucial requirement of large-scale releases. To effectively reduce malaria incidence, which is largely a rural disease, the high rate of male mosquito releases together with the associated cost of mass production make the implementation of such technologies challenging, especially in extensive areas of sub-Saharan Africa.

To overcome this, Burt and Deredec suggested the design of a self-limiting genetic tool, termed Y chromosome-linked genome editor (YLE)^22^, that would in theory have a more prolonged suppression effect in the target insect population. This technology is based on a genetic construct located on the Y chromosome that creates mutations that induce sterility or lethality when inherited by the female offspring but have no effect in the originator male or the male offspring (Fig. 1a). The genetic element encoding the editor would not increase its frequency in the population, but assuming no fitness costs, it is expected to remain at the released frequency because it follows Mendelian inheritance and is not selected against: the YLE construct only causes harm in the female offspring, where the Y chromosome is not present, and there is no alternative allele or chromosome in the YLE-bearing males to compete with. This lack of selection against the YLE has a pivotal consequence: if releases of males carrying the transgene are halted, the population would not immediately recover (as occurs with many other self-limiting strategies such as SIT, RIDL, or a female-specific RIDL), but the level of initial suppression would remain indefinitely^22^ (Fig. 1b).

**Fig. 1:**
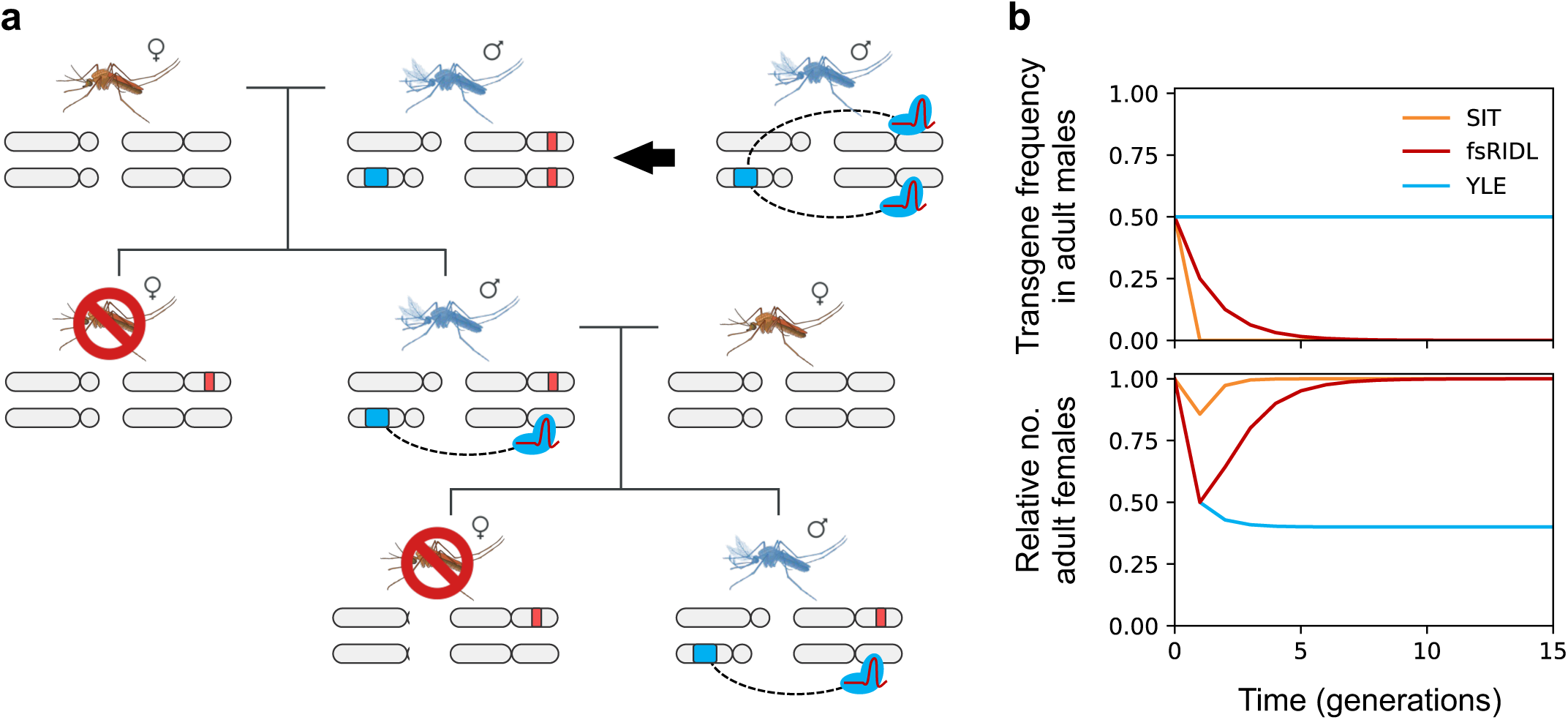
Dynamics of the Y chromosome-linked genome editor strategy. (**a**) Schematic representation of a Y chromosome-linked genome editor construct (in blue) that generates dominant mutations in genes required for female survival/fertility (in red) in the male germline through nuclease activity. These mutations are detrimental when inherited by females (causing lethality or sterility) but do not affect males. Because the genetic construct is located on the Y chromosome, which is always inherited by males, all the males in the offspring inherit it, allowing this process to be maintained through generations. (**b**) Modelling of the transgene frequency in adult males (top) and relative number of adult females (bottom) in a target mosquito population upon a single male release equivalent to 50% of the initial male population for SIT (orange), fs-RIDL (red) and YLE (blue).

These features make YLE technologies of great interest; however, the development of genetic strategies to control *Anopheles* mosquitoes that rely on the Y chromosome have proven to be challenging. This is mainly due to the highly heterochromatic nature of this chromosome and to the abundance of repetitive sequences that have prevented a complete chromosome assembly, hampering precise manipulation and site-specific transgene integration^23–25^. Furthermore, transgenes inserted on the Y chromosome of *Anopheles* species exhibit silencing due to heterochromatin and variable levels of expression due to position-effect variegation (PEV), as it occurs in *Drosophila*^23,26,27^. Besides these difficulties, meiotic sex chromosome inactivation (MSCI) is a major barrier to expression of Y-linked genes in the germline, which is a necessary pre-requisite for a YLE^28–30^. Nonetheless, germline expression of Y chromosome-linked transgenes has been previously achieved in *An. gambiae* using transcriptional control sequences that are activated prior to meiosis^23,31^.

Here, we used this knowledge to create a Y chromosome-linked Cas9-based system that is active in the male germline of *An. gambiae* mosquitoes, and we isolated an 11-base pair deletion in the *doublesex* gene that produces dominant female-specific sterility. Finally, we demonstrate that this deletion can be homed at a high rate using the Y-linked Cas9. To our knowledge, this represents proof of principle for the first genetic control strategy linked to the Y chromosome for the control of *An. gambiae* populations, and mathematical modelling data show that this technology is considerably more efficient than other self-limiting approaches such as SIT and various transgenic versions thereof. Such YLE may achieve efficient local suppression with a low risk of invading neighbouring populations, thereby overcoming the potential regulatory hurdles that more invasive strategies might face. Thus, this technology could be used both as a confined method for vector control and as a middle step in a phased malaria control programme aiming at using self-sustaining strategies.

## RESULTS

### Y chromosome-linked Cas9 activity leads to high levels of homing

In the attempt to build a YLE technology, we first investigated Cas9 activity from the Y chromosome by designing a genetic construct (YLE^dsx^) containing a gene coding for the endonuclease Cas9 under the control of the germline promoter *vas2*, and a gRNA, ubiquitously expressed, targeting the female-specific exon of *doublesex* (*dsx*) (Fig. 2a, 2b). The choice of *dsx* was advantageous due to the fact that, among our transgenic mosquito colonies, we harboured a strain with a *dsx* null allele that is marked with the dominant fluorescent marker gene GFP (*dsxF^GFP-null^*)^8^. The nature of this allele removes the Cas9-gRNA target site so that, in presence on the YLE^dsx^ construct, Cas9 cleavage of the wild type allele in the homologous chromosome, and repair using the *dsxF^GFP-null^* allele as a template, would lead to homing of the latter; this would be easily detected as an increase in the inheritance of GFP in the progeny of YLE^dsx^ males (super Mendelian inheritance) (Fig. 2c). The YLE^dsx^ construct was integrated in the genome of two independently originated *An. gambiae* strains, YLE^dsx^-A and YLE^dsx^-B, that each harboured ‘docking’ sites for site-specific recombination at different locations on the Y chromosome. We observed rates of inheritance of the *dsxF^GFP-null^* allele much higher than that expected from Mendelian inheritance, in both YLE^dsx^-A (81.5% inheritance) and YLE^dsx^-B (92.8% inheritance) backgrounds (Fig. 2d), indicative of high levels of Y chromosome-linked nuclease activity in the germline. While the results obtained for the two strains are not significantly different (P=0.0694; Mann-Whitney test), higher variability was observed in Cas9 activity for strain YLE^dsx^-A, suggesting that this locus might be more exposed to position-effect variegation and thus to a higher variability in transgene expression. Consistent with this hypothesis, in the offspring of YLE^dsx^ males with lower frequency of *dsxF^GFP-null^,* sequencing of the *dsx* target site showed very low level of mutations resulting from non-homologous end joining repair mechanisms, ruling out the possibility that reduced homing could have been the result of the emergence of alleles resistant to Cas9 cleavage^9,33^ (Supp. Table 1).

**Fig. 2:**
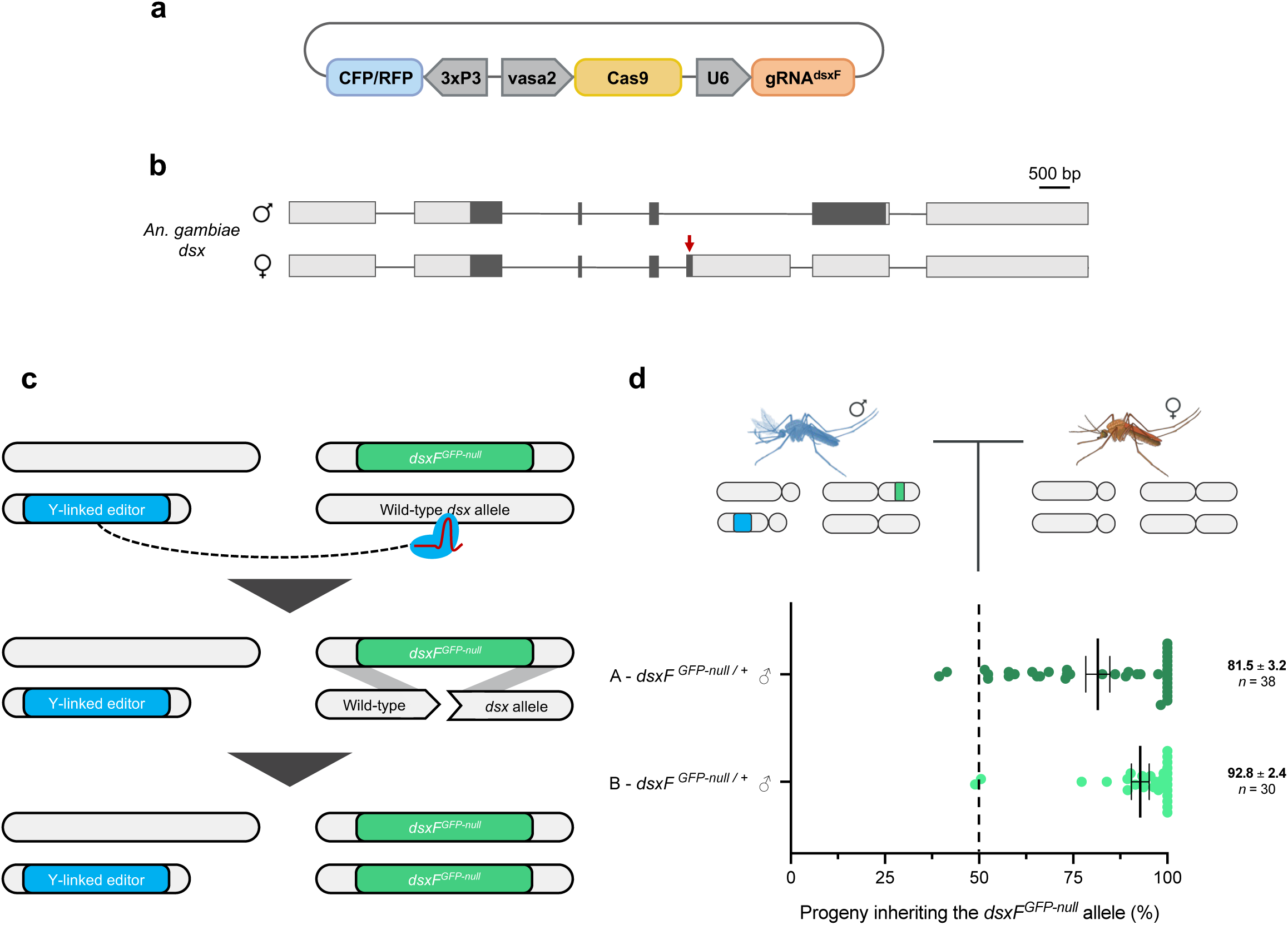
Generation of a Y chromosome-linked genome editor and evaluation of Y-linked Cas9 activity. (**a**) Genetic construct used to generate a YLE based on Cas9 expression under the germline *vasa2* promoter, targeting the female isoform of the *doublesex* gene. (**b**) Schematic representation of the *An. gambiae doublesex* gene, showing the male isoform (top) and the female isoform (bottom). Boxes represent exons and lines represent introns. CDS and UTRs are shown in dark and light colours, respectively. Introns are not drawn to scale. Red arrow indicates the site targeted by the YLE. (**c**) Illustration of homology-directed repair (homing) with a modified *dsx* allele (*dsxF^GFP-null^*) used to evaluate the Cas9 activity of the YLE^dsx^ strains. (**d**) Percentage of progeny from YLE^dsx^-A males (dark green) and YLE^dsx^-B males (light green) that inherited the *dsxF^GFP-null^* allele when crossed to wild-type individuals (n≥30). Each dot corresponds to the progeny of a different mating couple. Dashed line indicates the expected Mendelian inheritance. Numbers on the right indicate the mean and the s.e.m., and the size of each sample (n).

### An 11-base pair deletion in the coding sequence of *doublesex* causes female dominant sterility

When carrying out routine mosquito husbandry activities, in the offspring of YLE^dsx^ transgenic males, we observed a high frequency of genetic females showing an intersex phenotype resembling that found in females homozygous for the null mutation in the female-specific isoform of *dsx* (*dsxF^GFP-null^*)^8^. However, in our experimental design, Cas9-induced mutations generated in the reproductive organs of YLE^dsx^ males were expected to be present in heterozygosity in the progeny. Based on the knowledge that in *An. gambiae dsx* is a haplo-sufficient gene and that the disruption of the female-specific exon is recessive, we considered two most likely scenarios to explain the phenotype retrieved in our experiments: 1) conversion of both *dsx* wild-type alleles into a null mutation could have occurred in females with an intersex phenotype as a result of both germline activity of Cas9 in the transgenic male progenitors and paternally deposited nuclease; or, 2) based on the knowledge that sex-specific dominant negative mutations in the *dsx* gene have been identified in *Drosophila*^34–36^, a dominant negative mutation in the *dsx* gene was generated that is able to produce an intersex phenotype in females when in heterozygosity. Consistent with the second hypothesis, amplicon sequencing of the *dsx* target locus in pools of genotypic females showing the intersex phenotype, and Sanger sequencing in single individuals (over 400 individuals in total) revealed the presence in heterozygosity of an 11-base pair deletion within the coding sequence of the *dsx* female-specific exon (referred to as *dsxF^D11^* hereafter) (Fig. 3a). By contrast, amplicon sequencing of the *dsx* locus in pools of females retrieved from the same experimental batch and showing a wild-type phenotype (over 200 individuals in total) revealed the presence of this mutation only in 4 individuals. The low frequency at which this deletion is found in this pool of females may indicate either a non-complete penetrance of the *dsxF^D11^* mutation or a misidentification at pupal stage of these individuals when the phenotype was classified. Next, we conducted further analysis to explore the phenotype of intersex females. Dissections of 15 adult individuals revealed the presence of male-specific traits including incompletely formed claspers and organs resembling male accessory glands in conjunction with female-like antennae and apparently normal ovaries and spermathecae (Supp Fig. 1). Most importantly, a phenotypic assay was conducted to assess the fertility of intersex females. After mating and providing a blood meal, intersex females were unable to lay eggs, confirming full sterility for these individuals (Fig. 3b).

**Fig. 3:**
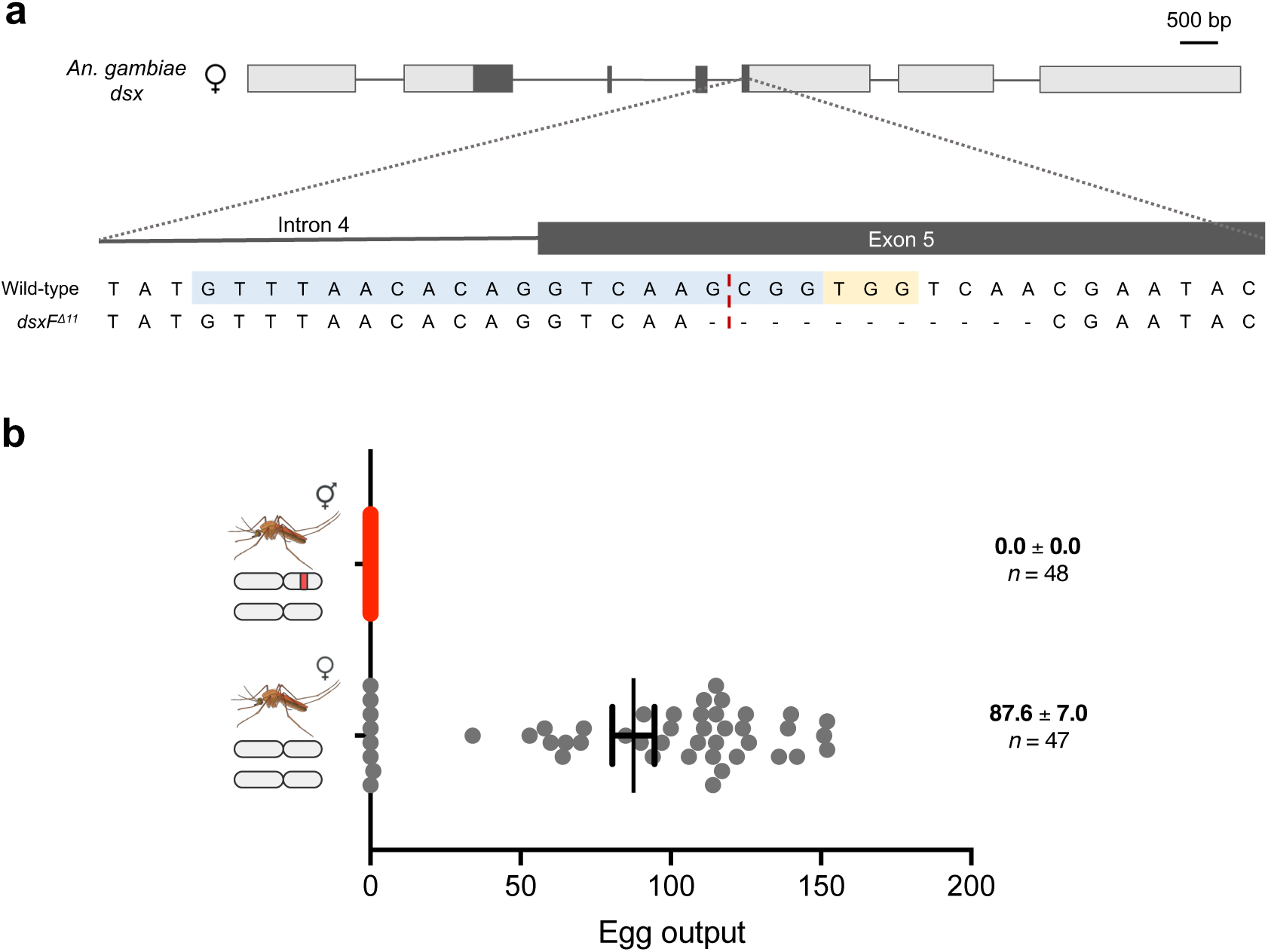
The *dsxF^Δ11^* allele and its dominant effect in the fertility of female individuals. (**a**) Intron 4-exon 5 boundary in the *dsx* gene. The gRNA sequence is highlighted in light blue and the PAM of the target site is shown in light yellow. The expected cut site is represented by a red dashed line. The sequence at the bottom shows the female-specific sterility-causing *dsxF^Δ11^* mutation. (**b**) Females with an intersex phenotype bearing a copy of the *dsxF^Δ11^* allele and females with a wild-type phenotype were mated with wild-type males and subsequently were given access to a blood meal. Each dot represents the progeny of each female. Numbers on the right show the mean and the s.e.m., and the size of each sample (n).

### Building a YLE strategy that biases the inheritance of the female-specific dominant mutation in the progeny

The existence of female-specific dominant negative mutations at the *dsxF* locus in *An. gambiae* is especially relevant for the strategy described here because it allows the possibility to use *dsx* as a target of a YLE to induce sterility in the female offspring. Because the *dsxF^D11^* allele is unrecognisable by the gRNA present in the YLE^dsx^ construct, this dominant mutation could be integrated in a system that biases the inheritance of *dsxF^D11^* by inducing cleavage of the wild type allele on the homologous autosome, from a source of Cas9 on the Y chromosome, and its conversion, through homology-directed repair, to the *dsxF^D11^* allele (‘homing’). We found that the *dsxF^D11^*allele was inherited by an average of 72.7% and 94.5% in the offspring of YLE^dsx^-A and YLE^dsx^-B males respectively (Fig. 4a). Consistently, females inheriting this mutation showed the intersex phenotype (see Supp. Fig. 1). New mutations at the *dsx* locus were found at very low frequencies (at an average of 1% and 3% of the individuals in the progenies of YLE^dsx^-A and YLE^dsx^-B males respectively), similarly to what we observed in the experiments conducted using the *dsxF^GFP-null^*allele (Supp. Tables 2-3). In one of these sets of progenies, in which all the sibling females showed an intersex phenotype, a one-base pair deletion within the coding sequence of exon 5 was found in heterozygosity in 28.36% of the individuals, indicating that this is likely another dominant mutation (Supp. Table 3, Supp. Fig. 2).

**Fig. 4:**
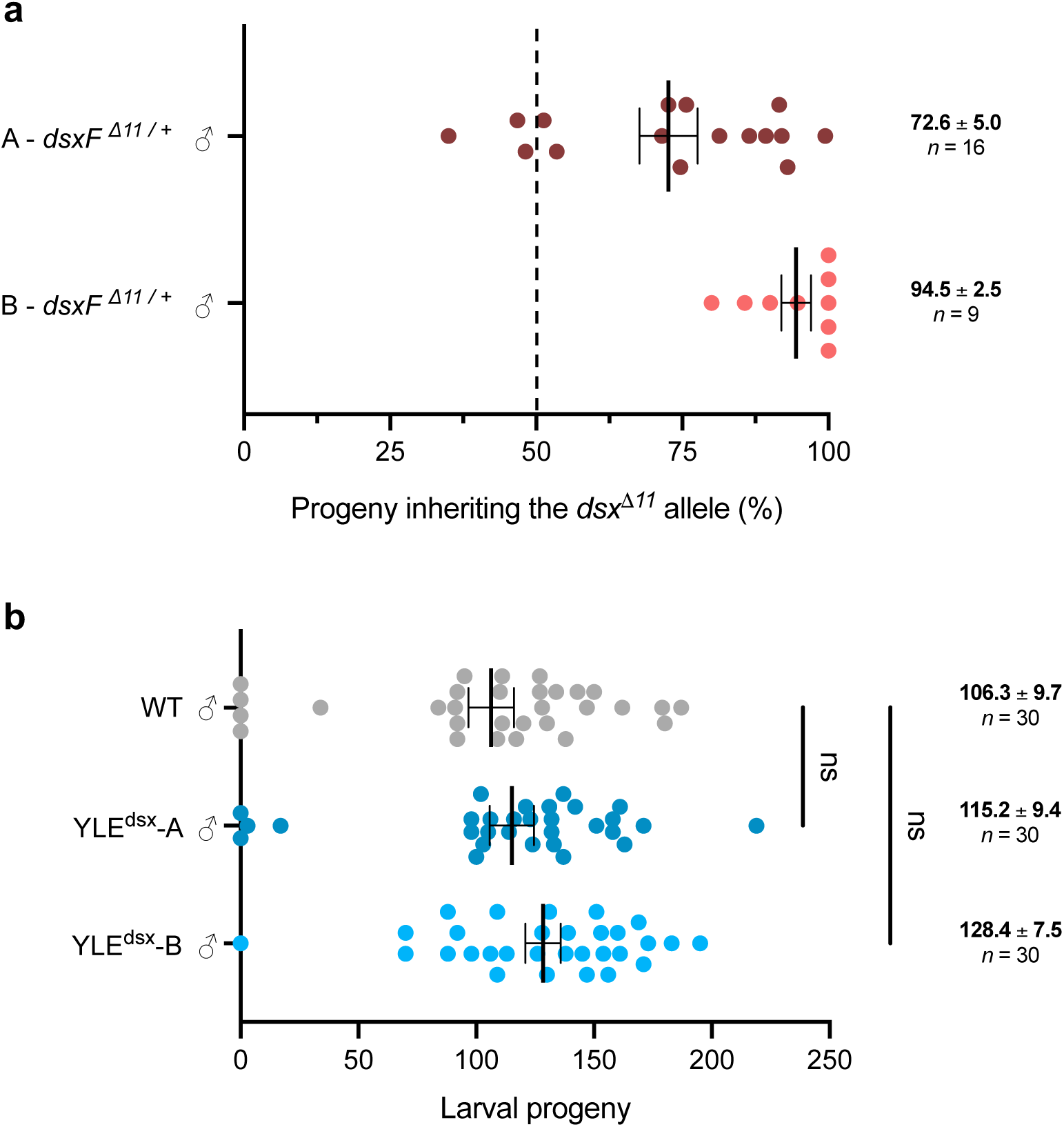
Transmission rate of the *dsxF^Δ11^* allele and fertility of YLE^dsx^ males. (**a**) Percentage of progeny from YLE^dsx^-A males (maroon) and YLE^dsx^-B males (red) that inherited the *dsxF^Δ11^* allele when mated with wild-type individuals. Each dot corresponds to the progeny of a different mating couple. Dashed line indicates the expected Mendelian inheritance. Numbers on the right indicate the mean and the s.e.m., and the size of each sample (n). (**b**) YLE^dsx^ males from both strains (A and B) were mated with wild-type females. A Kruskal-Wallis test revealed no significant differences (P = 0.2322) in the number of larvae produced by females mated with YLE^dsx^ males and those mated with wild-type males. Each dot represents the progeny of each female. Numbers on the right-hand side show the mean and the s.e.m., and the size of each sample (n).

### The YLE^dsx^ transgene does not pose a fertility cost on male mosquitoes

The YLE^dsx^ strains described in this study produce progeny consisting of sterile females and transgene-carrying males. To be most useful as a genetic control strain, males bearing a YLE construct should have comparable fitness to wild-type males, since the persistence of suppression is directly related to this parameter^22^. Wild-type females mated with YLE^dsx^-A and YLE^dsx^-B males (n = 30) produced an equivalent number of larvae to females mated with wild-type males, hence providing evidence that, under standard insectary conditions, the fertility of YLE^dsx^-A and YLE^dsx^-B males is not significantly different from the fertility of wild-type males from the same colony (Fig. 4b).

### Mathematical modelling suggests a great potential for YLEs as self-limiting genetic control tool

To assess the potential of the YLE^dsx^-B strain as a tool to suppress a target mosquito population in the wild, we used the life history traits observed in laboratory conditions for this strain as parameters in a mathematical model. In this simulation, released transgenic males harbour both the YLE^dsx^ construct and a female-specific dominant negative mutation, causing females sterility, with total penetrance; an equivalent scenario to the release of YLE^dsx^ males heterozygous for the *dsxF^D11^* allele described in this study (YLE^dsx^; *dsxF^D11/+^*). Two parameters are unknown in this model: 1) the rate of *de novo* mutations in that fraction of YLE^dsx^ males that, in subsequent generations post-release, may have two unmodified *dsx* alleles; and 2) the landscape of other mutations in that locus that cause an intersex phenotype, and their penetrance. For these reasons, a conservative approach was taken with the assumption that every male that did not inherit the released dominant mutation would generate a recessive one that would be transmitted to their progeny at the same frequency as the dominant mutation in released males.

We used this model to determine the release rate required to achieve 95 and 99% suppression in 36 generations for a target mosquito population with an intrinsic rate of increase of R_m_=6, assuming equal sized releases in every generation. The model showed that the developed YLE^dsx^ would require 7-8 times lower releases than an optimal SIT strategy, 3-4 times lower releases than optimal RIDL or fs-RIDL technologies, and 2-3 times lower releases than an X-shredder (Table 1). Nevertheless, our YLE^dsx^ strain would still need 4-5 times larger releases than an optimal YLE. Original mathematical modelling on this strategy showed that the release rates could be reduced if YLE males also carry an autosomal X-shredder construct that leads to a disproportionate transmission of the Y chromosome^22^. Additionally, our model with the YLE^dsx^ strain parameters shows that the combination with an autosomal X-shredder would halve the release rates required (Table 1).

**Table 1.** Comparison of the release rates required to suppress a target mosquito population with YLE^dsx^ males and other optimal self-limiting technologies. Release rates required to achieve a suppression level of 95 and 99% in a target mosquito population with an intrinsic rate of increase (*R_m_*) of 6, assuming releases for 36 generations. Release rates were calculated as a proportion of the initial number of males in the target population, for the developed YLE^dsx^-B strain, an optimal YLE causing female-specific sterility, an optimal SIT, an idealised RIDL (causing unspecific lethality after density-dependence), an optimal fs-RIDL (causing female-specific lethality after density-dependence), an idealised autosomal X-shredder, and males from the YLE^dsx^-B strain bearing an autosomal X-shredder construct in heterozygosity (+/-) or in homozygosity (+/+). For the optimal autosomal approaches, it is assumed the release of homozygous males. Numbers in square brackets indicate how many times larger the release rate would have to be compared to the release of males from the YLE^dsx^-B strain to achieve the same level of suppression.

| Suppression level (%) | YLE<sup>dsx</sup>-B | Optimal YLE | Optimal SIT | Optimal RIDL | Optimal fs-RIDL | Optimal X-shredder | YLE<sup>dsx</sup>-B ; X-shredder (+/-) | YLE<sup>dsx</sup>-B ; X-shredder (+/+) |
| --- | --- | --- | --- | --- | --- | --- | --- | --- |
| 95 | 0.171 | 0.037<br>[0.2] | 1.297<br>[7.6] | 0.438<br>[2.6] | 0.668<br>[3.9] | 0.396<br>[2.3] | 0.114<br>[0.7] | 0.088<br>[0.5] |
| 99 | 0.183 | 0.041<br>[0.2] | 1.300<br>[7.1] | 0.440<br>[2.4] | 0.678<br>[3.7] | 0.402<br>[2.2] | 0.122<br>[0.7] | 0.095<br>[0.5] |

To understand the source of the difference between our YLE^dsx^ strain and an optimal YLE, we performed a sensitivity analysis of how the release rate required varies as a function of changes in the underlying genetic variables (Fig. 5). Among all the analysed variables, it is the rare production of recessive alleles that is the most critical, while the remaining parameters of the YLE^dsx^ are very close to the optimum values.

**Fig. 5:**
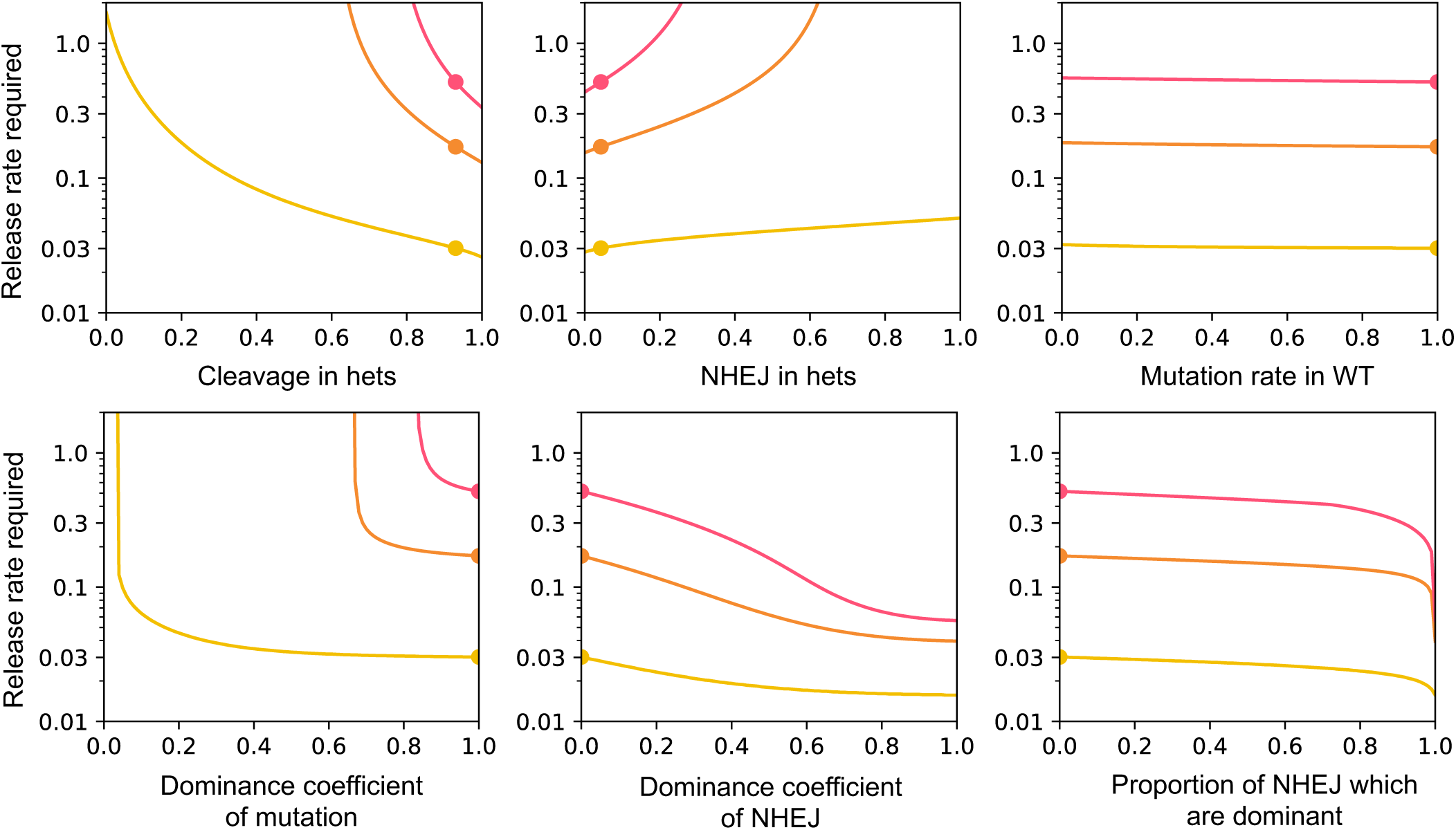
Release rates required to achieve 95% suppression of a target population within 36 generations with males from the YLE^dsx^-B strain as a function of various parameters, for *R_m_*= 2, 6, and 12 (yellow, orange, and red, respectively) Lines illustrate the release rates required as a function of each of the parameters, where all other parameters are set to baseline values (Supp. Table 4). Dots indicate the current value of the developed YLE^dsx^ for each parameter. *R_m_* is defined as the intrinsic rate of increase of the population (expected number of daughters produced by a female in the absence of density-dependent mortality). Analysed parameters shown in the top row are (left to right): cleavage rate in YLE males heterozygous for a mutation (i.e., probability of cleaving the unmodified target in YLE males bearing one mutated copy), probability of generating a new mutation in the target by non-homologous end-joining (NHEJ) given cleavage in YLE males heterozygous for a mutation, and probability of creating a mutation in the target site in YLE males that are wild-type homozygous. In the bottom row, analysed aspects include (left to right): dominance coefficient of the released mutation (i.e., if the mutation carried by released YLEdsx males is fully penetrant and always causes sterility in females, the dominance coefficient will be 1), dominance coefficient of new dominant mutations generated by NHEJ, and proportion of new mutations generated by NHEJ that are dominant.

Recessive mutations produced by non-homologous end joining repair mechanisms would accumulate in the target population because they could not be recognized and cleaved by the Cas9-gRNA complex and they would not be selected out as efficiently as dominant mutations, since females that are heterozygous for these alleles would be fertile and would contribute to their transmission. Nonetheless, this would not prevent a prolonged suppression effect in the target population following a single release of YLE^dsx^; *dsxF^D11/+^* males or after halting generational releases, nor the virtual suppression of a target mosquito population if releases continue (Fig. 6).

**Fig. 6:**
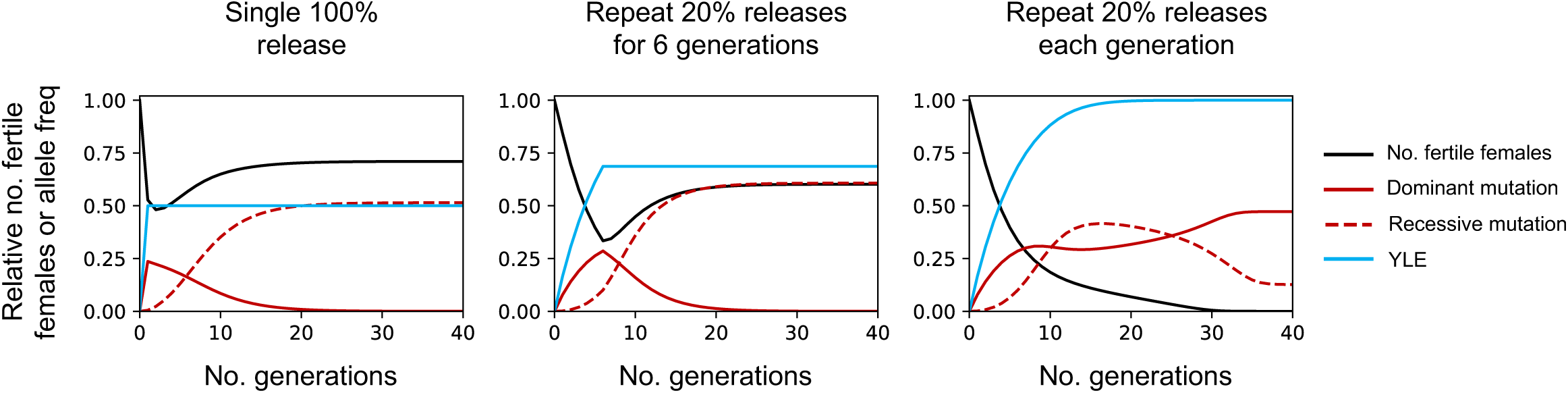
Time course of a population suppression upon releases of YLE^dsx^ males heterozygous for the dominant *dsxF^Δ11^* allele. The black line shows the relative number of fertile females in a target population across generations following a single release of YLE^dsx^; *dsxF^Δ11/+^* males equivalent to the initial number of males in the population (*left*), and releases of YLE^dsx^; *dsxF^Δ11/+^* males equivalent to 20% of the initial male population for 6 generations (*middle*) or indefinitely (*right*). The frequency of males bearing the YLE construct is represented in blue. The frequency of the released dominant *dsxF^Δ11^* mutation in the population is shown by the continuous red line, and the dashed red line indicates the frequency of emerging recessive mutations. *R_m_* = 6.

## DISCUSSION

In this study, we used the *vasa* regulatory sequences to drive the expression of a Y-linked Cas9 cassette in the germline of *An. gambiae*. We show high activity of the endonuclease that was able to promote efficient homing of a newly discovered female-specific dominant mutation (*dsxF^D11^*) at the *dsx* locus, ensuring that a high proportion of females inherit a copy of this mutation and are fully sterile. The differences observed at two distinct Y chromosome integration sites, in terms of overall Cas9 activity and variability in this activity among males with the same engineered Y chromosome, seem to indicate that this chromosome is affected by position-effect variegation through its length. This is particularly relevant for the development of future genetic control strategies linked to the Y chromosome in *An. gambiae*, and efforts to locate the relative positions of the two Y chromosome integrations described in this study could provide useful information for the identification of chromosome regions that are more transcriptionally active and less prone to silencing.

In addition, while mating competitiveness of YLE^dsx^ *versus* wild-type males was not assessed due to the constraints of small cage experiments, we observed that the fertility of these males was not significantly different. This would be especially pertinent in a release scenario of YLE males, as the transgene persistence in the wild and its suppression effect in the population over generations would be affected by potential fitness costs associated to the YLE construct.

The modelling for the YLE^dsx^-B strain showed that this self-limiting strategy could lead to significant reductions in the size of the target mosquito population, and the long-lasting effect of the YLE in the population after the release would allow for a phased suppression approach. On this premise, if YLE males were to be released at a 10% frequency of the initial male population, half of their offspring (i.e., the female portion) would be sterile each generation, resulting in a persistent 5% reduction in the population. If the same number of initial YLE males were to be released in a subsequent generation, it would represent a higher overall frequency and, in a rachet-like manner, cause a further decrease in the total population due to a higher proportion of females being eliminated. The described model demonstrated that generational releases of YLE^dsx^; *dsxF^D11/+^* males could potentially lead to the virtual collapse of the target population. While the release rates required for YLE^dsx^-B are four to five times larger than for the optimal YLE, these are still significantly lower than the release rates for other optimal self-limiting strategies. Given the cost and logistic challenges associated with mass rearing of mosquitoes, the use of YLE could provide substantial operational and resource efficiencies over current approaches to field releases of vector control agents. Furthermore, combining the YLE^dsx^ with an autosomal X-shredder would reduce the required release rates below those of an optimal YLE.

While sex-specific dominant mutations have been widely studied in the *doublesex* gene of *D. melanogaster*^35–37^, this was not the case in *An. gambiae* mosquitoes. The identification of the dominance of the *dsxF^D11^* allele is key to the YLE approach described here because it induces sterility in females. This dominant mutation carries an out-of-frame 11-base pair deletion within the coding sequence of the female-specific exon and introduces an early stop codon that would lead to the production of a truncated protein, which would be missing nearly all the highly conserved female-specific region of the DSX protein and would show a severely affected predicted structure^38^ (this would also be the case for the one-base pair deletion found to be dominant) (Supp. Fig. 2, 3). Given that the truncated protein would keep the DNA-binding and dimerization domain intact, the dominant phenotype might be explained by its interference with the wild-type DSX protein. This mechanism would also explain the observed differences in the external phenotype since the level of interference would be a stochastic process and the phenotype would vary among females. If this is the case, other mutations within the coding sequence of the *dsx* female-specific exon might have a dominant effect, as suggested by the finding of the dominant one-base pair deletion. Nonetheless, the *dsx^GFP-null^* allele in *An. gambiae* and other examples in *D. melanogaster* show that certain mutations within this region might be recessive^8,34,39^.

Mathematical modelling indicates that amongst the potential mutations in the target gene within the offspring that do not receive a homed allele, the relative ratio of mutations that are novel dominant negative *versus* recessive loss of function will significantly influence the efficiency of the YLE strategy. In an ideal scenario, 100% homing rates of the *dsxF^D11^*allele would solve this issue; however, this is unlikely to be achieved. To address this, target genes for this type of YLE should not easily tolerate modifications that include recessive mutations or alleles resistant to cleavage that restore the gene function. Although *dsx* is a functionally constrained gene, it would be worthwhile to explore the potential for genetic variation within the female-specific exon and its surroundings to generate dominant mutations over recessive or functional resistant ones.

Given the challenge of achieving 100% homing rates, a strategy based on multiplexing might be considered. While homing-based technologies generally aim to reduce or delay the emergence of resistance by targeting two different gRNA sites within the same gene, in this YLE strategy, multiple gRNAs to home a dominant mutation should also be designed to ensure that any repair outcome would result in a dominant effect. Alternatively, two different genes that can induce a female-specific dominant effect could be targeted. Additional targets could be explored by looking for genes which proteins form oligomers and have an essential role only in females, such as the *Act4* gene in *Aedes aegypti* and *Culex quinquefasciatus* mosquitoes, for which different in-frame deletions cause a dominant flightless phenotype in females^40,41^. Genes that are located in the X chromosome might also be used as targets, as long as they are not required in male spermatogenesis, given that the X chromosome will always be inherited by the female offspring. Since males only bear one copy of the X chromosome, the YLE could not rely on the homing of a dominant mutation but rather ensure that every generation a dominant negative allele is generated. Such a target might be found in the ortholog of the *D. melanogaster* gene *wupA*, which has been shown to induce embryonic female lethality when targeted with an endonuclease in the *D. melanogaster* male germline, suggesting that it is an haploinsufficient gene^42,43^.

With regards to preparatory work for its application, a YLE strain would be easily maintained and scaled up prior to field releases. In an insectary facility, male larvae can be automatically sorted due to the Y chromosome-linked fluorescent marker, and then crossed to wild-type females. If the aim is to release the YLE paired with a dominant mutation such as *dsxF^D11^*, given that it is not possible to generate a homozygous strain, it might be desirable to link this allele to a fluorescent marker in a way that it does not affect the dominant phenotype (e.g., by placing the marker in a nearby intron), and therefore ensure that all the released males bear this mutation. High-throughput larval screening using flow cytometry^44^ could allow effective selection of larvae containing the YLE construct and the dominant mutation(s).

The YLE^dsx^ strain presented here is a self-limiting genetic control tool for *An. gambiae* that would seem to be more effective than other self-limiting vector control technologies: it has a stronger effect on population suppression that is maintained for a longer period of time. At the same time, it is less invasive than self-sustaining approaches and it is expected to be more geographically confined. This YLE technology expands the toolbox of genetic control strategies and offers an option halfway between previously existing self-limiting technologies and self-sustaining ones.

## METHODS

### Plasmids used for the generation of the YLE strains

The plasmid p16510, which had been previously engineered^8^, contains a human-codon-optimized *Cas9* coding sequence under the control of the *vas2* promoter^31^, a gRNA targeting the intron 4 – exon 5 boundary of the *dsx* gene under the ubiquitous *U6* promoter, and a gene coding for *RFP* under the *3xP3* promoter. Additionally, a modification of this plasmid existed with the sole difference of a gene coding for *CFP* instead of *RFP*.

### Generation of the YLE strains targeting *dsx*

To generate the YLE^dsx^-A strain, embryos from the Y-AttP line^23^ were injected with solution containing the p16510-CFP plasmid (at 200 ng/μL) and a *vasa-integrase* helper plasmid (at 400 ng/μL) for integrase-mediated insertion. Microinjections were performed as described previously^45^. In order to create the YLE^dsx^-B strain, we injected embryos from the strain provided by Jaroslaw Krzywinski (bearing a transgene on the Y chromosome with attP sites for integrase-mediated recombination) with solution containing the p16510-RFP plasmid (at 200 ng/μL) and a *vasa-integrase* helper plasmid (at 400 ng/μL).

Surviving L1 larvae were screened for transient expression of the fluorescent marker in the anal papillae and parts of the ventral nerve chord. Larvae with and without this transient expression were reared separately and crossed to wild-type mosquitoes at adulthood. The progeny from these crosses was screened for expression of the fluorescent marker under the *3xP3* promoter, visible in the optic lobe of the head and the dorsal ganglia across the larval body. Successful integration events were detected by the exchange of the fluorescent marker in G1 male individuals that were RFP^-^ and CFP^+^ for YLE^dsx^-A, or GFP^-^ and RFP^+^ for YLE^dsx^-A.

### Assessment of homing levels of the *dsxF^GFP-null^* allele

To ensure that each progeny analysed for homing is coming from a different male, the experimental crosses were performed as described previously^46^ and explained below. YLE^dsx^ males were initially mated with females heterozygous for the *dsxF^GFP-null^* allele. Individuals in the progeny with expression of the marker linked to the YLE and GFP (linked to *dsxF^GFP-null^*) were selected. To ensure that YLE^dsx^ males used in the experimental cross would not have inherited other mutations in the target site that would prevent homing of *dsxF^GFP-null^*, the selected males were mated with wild-type females. Around 600 double-positive individuals were selected in the offspring at pupal stage and placed in a large cage. 100 females from the *An. gambiae* wild-type colony were sexed at pupal stage and placed in a separate cage. Six days after emergence, at the time of the programmed dusk, females were introduced into the male cage in groups of 6-8. Mating pairs falling to the bottom of the cage were isolated by gently placing a 25 mL plastic glass over them until copulation terminated. Once the couple separated, the plastic glass was brought over a thin piece of cardboard previously left on the bottom of the cage and the mated mosquitoes were transferred to another cage. Mated females were kept in this separate cage and males were removed to avoid a new mating with another female. 50-60 mated females were captured. The following day to the mating, females were fed with blood and 3 days later they were allowed to lay eggs individually. Hatched L1 larvae were screened for the presence of GFP. Larvae from progenies with low homing rates of *dsxF^GFP-null^* were pooled to perform genomic DNA extraction (Wizard® Genomic DNA Purification Kit, Promega) and subsequently do amplicon sequencing on the *dsx* target to check for the presence of mutations that could have prevented the homing. Amplicon sequencing data were analysed using CRISPResso2^47^.

While the procedure to obtain YLE^dsx^; *dsxF^GFP-null/+^* males was the same for both YLE^dsx^ lines, the YLE^dsx^-B line was screened by eye under the fluorescence microscope, but the YLE^dsx^-A line was screened using the COPAS system to unambiguously detect males expressing GFP. Homing of the *dsxF^GFP-null^* allele in the YLE^dsx^-A line was assessed by eye under the fluorescence microscope and evaluated only in females to avoid confusion with CFP^+^ only males.

Statistical differences in homing rates between both strains were assessed using a Mann-Whitney test.

### Assessment of homing levels of the *dsxF^Δ11^* allele

To ensure that the YLE^dsx^ males used for the experimental cross had the markerless *dsxF^Δ11^* allele, the offspring of different females from these lines were reared separately. Individuals were sexed at pupal stage, looking for progenies where all the females showed the intersex phenotype (indicating that all of them had the dominant mutation, and therefore their sibling males did too). The males of those progenies were selected to perform the experiment: 50 YLE^dsx^ males supposedly bearing the dominant mutation were allowed to mate for 5 days to 60 wild-type females in a cage. Females were then fed blood and allowed to lay eggs individually 3 days later. Progenies were raised separately and sexed at pupal stage. Females from each progeny were then pooled to perform genomic DNA extraction (Wizard® Genomic DNA Purification Kit, Promega) and subsequently do amplicon sequencing on the *dsx* target to check the frequency of the *dsxF^Δ11^* allele in the offspring. Amplicon sequencing data were analysed using CRISPResso2^47^.

### Fertility assays

YLE^dsx^ males from both lines were crossed with wild-type females in groups of 50. They were left to mate for 5-7 days before being blood fed. A minimum of 30 females were allowed to lay eggs individually by placing them in separate cups 3 days after blood feeding. After 2 nights, the number of eggs laid by each female was recorded. The number of larvae hatching from each progeny in the following days was recorded as well.

Females with an intersex phenotype were crossed with wild-type males in groups of 60. They were left to mate for 5-7 days before being blood fed. A minimum of 50 females were allowed to lay eggs individually by placing them in separate cups 3 days after blood feeding. After 2 nights, the number of eggs laid by each female was recorded. The number of larvae hatching from each progeny in the following days was recorded as well. The same procedure was followed with wild-type females as a control.

Statistical differences against wild-type reference crosses were assessed using a Kruskal-Wallis test.

### Dissections of females showing an intersex phenotype

Adult females with an intersex phenotype were anaesthetised on ice for 5-10 minutes. Individuals were then place on a glass slide with PBS solution and were dissected under a stereomicroscope using two needles. One needle was used to hold the thorax of the females while the other one was placed on the last segment of the abdomen to pull. Gonads and spermathecae were isolated from the rest of the body for visualization and recording under an EVOS XL Core Cell imaging system. Dissections of wild-type males and females were also performed for comparison.

### Mathematical modelling

#### Overview

The model developed to simulate the YLE^dsx^ has a deterministic structure, based on the one developed by Burt and Deredec^22^. It simulates a single, infinite-sized population where mating occurs randomly, with discrete, non-overlapping generations and three life stages (larvae, pupae and adults). We first model a Y-linked locus with two alleles: the wild-type (*Y*) and the YLE (*y*); and an autosomal tharget locus with three alleles: the wild-type (*A*), a cleavage-resistant recessive mutation (a) and dominant mutation (*α*). No functional cleavage-resistant alleles were considered. Therefore, considering both the Y-linked and autosomal loci, there are a total of 12 male genotypes and 6 female genotypes. We then extend the model to simulate the YLE in the presence of an autosomal sex distorter construct which results in cleavage of the X chromosome during spermatogenesis, increasing the proportion of Y-bearing sperm (X-shredder). We include a second autosomal locus with two alleles, the wild-type (*B*) and the X-shredder construct (*b*). For the extended model, considering the Y-linked and both autosomal loci, there are a total of 90 male genotypes and 45 female genotypes. Using these models it is also possible to simulate a range of comparative strategies including release of sterile males (SIT), release of individuals carrying a mutation which causes death after density dependent mortality in both sexes (RIDL) or only in females (fsRIDL) and an X-shredder alone. All simulations were performed in Julia, a scientific programming language^48^ within the Jupyter notebook interface^49^. Code used for the simulations is available on GitHub (https://github.com/KatieWillis/YLESimulator).

#### Population biology

To simulate a population through time in each generation females produce *f* eggs which are fertilised by males assuming random mating and that the number of males is not limiting to the number of eggs fertilised. Larval survival to the pupal stage is density-dependent according to the Beverton-Holt model, where the probability of survival is 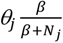, *θ_j_* is the density-independent probability that a juvenile survives to adulthood, *β* determines the strength of density-dependent mortality and *N_j_* is the total number of juveniles in the population. In this study all results are reported as relative population sizes of biting females compared to the starting population at equilibrium, therefor they are not affected by the value of *β*. The intrinsic rate of increase (*R_m_*) of the wild-type population is 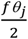. A value of 6 was used for the *R_m_* in all simulations unless otherwise stated^50^.

#### Gamete transmission

In males carrying the YLE, the probability of gamete transmission can be altered due to cleavage of the WT autosomal target. Cleavage of the WT allele occurs with probability *c*, and, in WT heterozygotes repair of the cleaved chromosome occurs by non-homologous end-joining with probability *j*, converting the WT *A* allele to a cleavage-resistant allele, a or *α*, with probability *p* and 1 − *p* respectively. Alternatively, the cleaved chromosome is repaired through homology-directed repair, converting the *A* allele to either an a or *α* allele, depending on the alternative allele present in the genotype. In males carrying the YLE and two copies of the WT autosomal allele, cleavage and mutation of each WT allele occurs with probability *μ*, converting the *A* allele to a cleavage-resistant allele, a or *α*, with probability *q* and 1 − *q* respectively. Since the editor is located on the Y chromosome, there is no change in gamete transmission due to YLE activity in females. Haploid gamete genotypes are generated assuming random segregation of the edited (or unedited) parental diploid genotype, and the proportion of zygotes of each diploid genotype is calculated assuming random pairing of of male and female gametes. To model the effects of the autosomal X-shredder, males carrying at least one *b* allele produce Y and X bearing sperm at a ratio of *m* : 1 − *m*. When *m* is equal to 1 all gametes produced by males carrying the *b* allele contain a Y chromosome (i.e. all offspring fertilised by these sperm are male) whereas when *m* is equal to 0.5 there is no sex-ratio distortion. Descriptions of all inheritance parameters and their values when modelling the YLE^dsx^ can be found in Supp. Table 4. Note that for SIT, RIDL, fsRIDL, and the X-shredder alone no editing occurs, therefore *c*, *j*, μ, *p* and *q* were equal to 0. When modelling strategies with an idealised autosomal X-shredder *m* was equal to 1.

#### Fitness effects

The fitness of individuals can be affected by the presence of alleles which disrupt the function of the *A* locus. To model this the fitness of *A*/*A* individuals is normalised to 1; and the relative fitness of genotypes homozygous for the dominant mutation *α*/*α* is 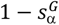; homozygous for the recessive mutation a/a is 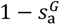; heterozygous for the dominant mutation and WT *A*/*α* is 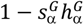; heterozygous for the recessive mutation and WT *A*/a is 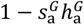; and heterozygous for the dominant and recessive mutation a/*α* is 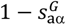. Here, 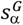, 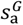, and 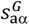 are selection coefficients, and 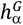 and 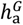 are dominance coefficients, each of which were allowed to vary depending on sex (*G*). Parameterisation in this way allows for modelling of female-specific fitness effects of the YLE and comparator strategies such as the fsRIDL, as well as bi-sex fitness effects such as for in SIT and RIDL. For simplicity, it was assumed that the A locus is essential for survival to reproductive maturity. Depending on the strategy being modelled, the genotype-dependent fitness costs are either applied before density-dependent mortality (as if larvae die before reaching pupation, as in SIT) or after (as if pupae die before maturing into reproductively active adults, as in RIDL, fsRIDL and YLE). In the case of female-specific costs acting after density-dependent mortality, this is equivalent to females being both infertile and unable to bite.

Small and large cage trials of *An. gambiae* males carrying an X-shredder revealed two fitness effects of the X-shredder: reduced fertility in males carrying an X-shredder and reduced survival of daughters with X-shredder fathers due to inheriting X chromosomes exposed to the X-shredder^10,51^. For simplicity, we incorporate only the first cost since it effects all X-shredder males and is expected to be the most impactful, whereas the second cost is restricted to the rare females produced from X-shredder males. Since we assume random mating and that the number of males is not limiting to the number of eggs fertilised, reduced male fertility is modelled by i) reducing the survival of males from pupae to adult by a factor of 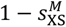, where 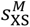 is the fitness cost of the X-shredder males and ii) reducing the number of eggs laid by all females by a factor of 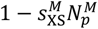, where 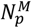 is the proportion of X-shredder carrying males in the population at the pupal stage. For simplicity we assume the fitness costs are the same whether males carry one or two X-shredder constructs. Descriptions of the fitness parameters and their values for each strategy can be found in Supp. Table 5.

#### Time series simulations and releases

When modelling the YLE^dsx^, released males were assumed to carry a YLE construct and a single copy of a dominant female-specific cleavage-resistant allele (*A*/*α*). In the case of SIT, RIDL, fsRIDL and the X-shredder alone, released males carried a WT Y-chromosome (*Y*) and were homozygous for the fitness inducing modification (*α*), the fitness effects of which varied depending on the strategy being modelled. The population was censused at the zygote stage where autosomal allele frequencies were calculated in males and females independently and averaged, whereas frequencies of Y-linked alleles refer only to males. Estimated release rates required were accurate to 3 decimal places.

## ACKNOWLEDGMENTS

We would like to thank Jaroslaw Krzywinski for providing one of the *An. gambiae* Y docking-site strains used in this work. We also thank John Connolly and the Target Malaria Regulatory Team for their valuable feedback. This work was supported by the Bill & Melinda Gates Foundation and Open Philanthropy.

## AUTHOR CONTRIBUTIONS

T.N. and A.C. conceived the project; I.T., T.N. and F.B. designed the research; I.T. and M.G. performed research; I.T., T.N. and F.B. analysed data; K.W. and A.B. performed mathematical modelling; and I.T., T.N. and F.B. wrote the paper with input from all authors.

