## Supplementary material for "A Y chromosome-linked genome editor for efficient population suppression in the malaria vector *Anopheles gambiae*"

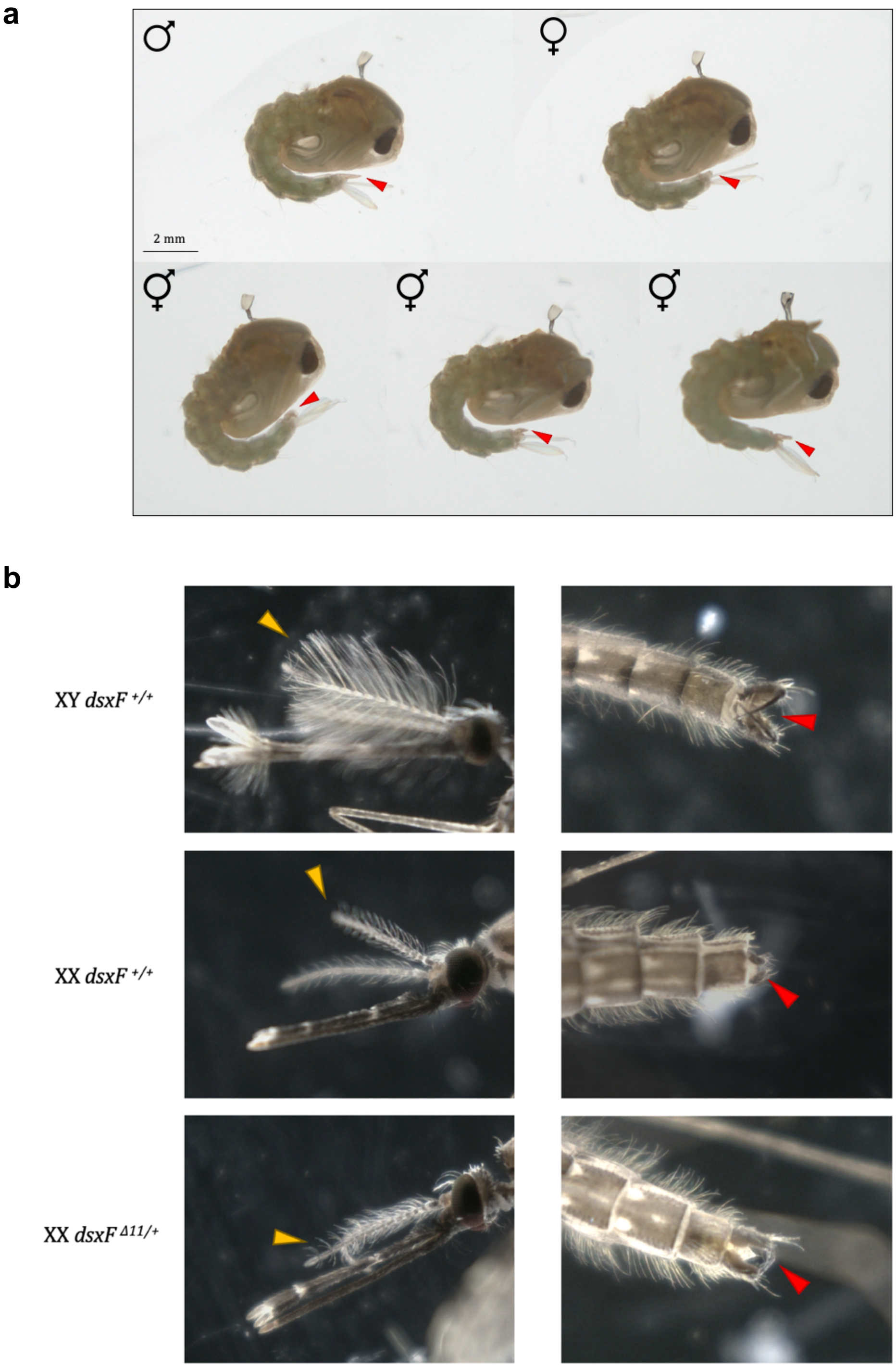


**Supplementary figure 1. Intersex phenotype at pupal and adult stages of chromosomally female individuals heterozygous for the *dsxF*$^{\Delta11}$ allele.** (**a**) Females bearing one copy of the *dsxF*$^{\Delta11}$ mutation showed an intersex phenotype at pupal stage (bottom row), which could be observed in the varied development of the genital lobe (red arrows), which was much more developed than in wild-type females (top-right) but less developed than in wild-type males (top-left). (**b**) At adulthood, females heterozygous for *dsxF*$^{\Delta11}$ (bottom) have female-like pilose antennae (yellow arrows) and a pair of claspers (red arrows) that are dorsally rotated in comparison to the claspers in wild-type males (top), which wild-type females lack (middle).

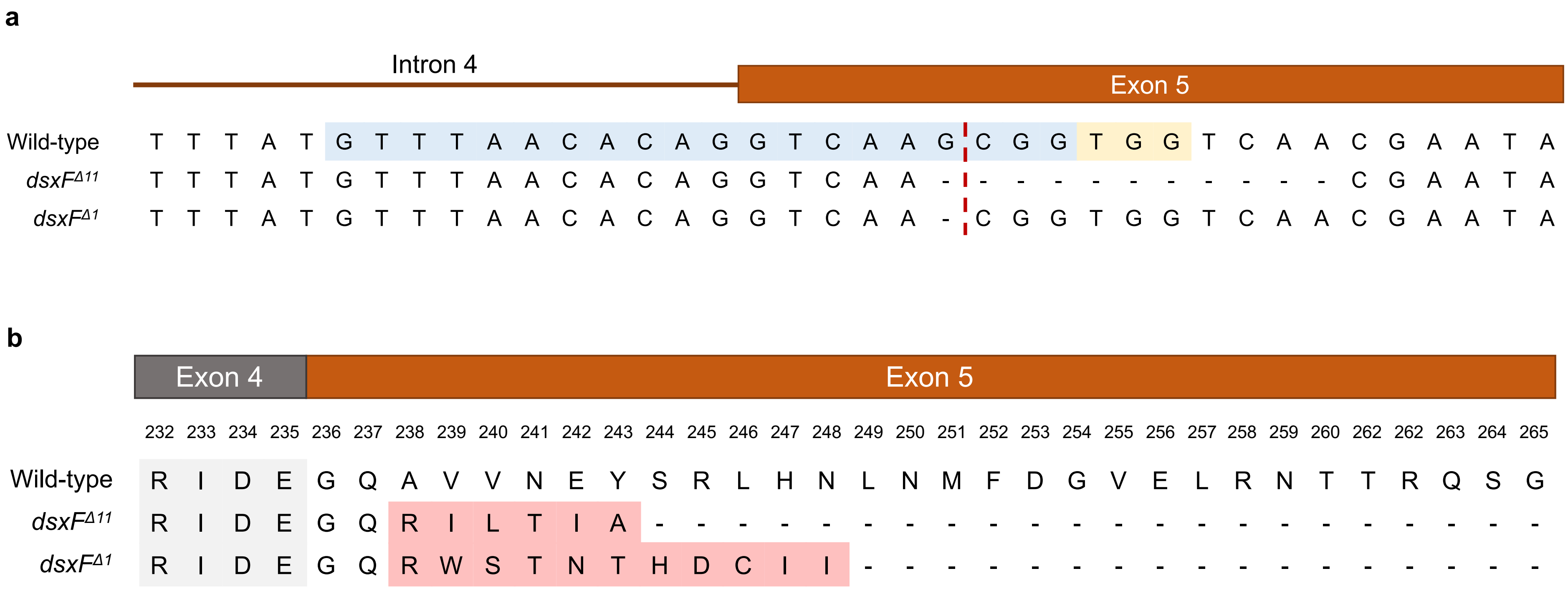


**Supplementary figure 2 – Comparison of the nucleotide and amino acid sequences of the two dominant negative mutations identified in the *dsx* locus of *An. gambiae*. (a)** Representation of the target sequence environment in the intron 4 – exon 5 boundary of the *dsx* gene. The gRNA (and target sequence) is highlighted in blue, the PAM at the target site is highlighted in yellow, and the cut site is shown by a dashed red line. The two dominant mutations identified (the 11bp deletion - *dsxF*$^{\Delta 11}$ - and the 1 bp deletion - *dsxF* $^{\Delta 1}$-). **(b)** Representation of the amino acid sequence encoded in the end of exon 4 and the coding sequence of exon 5 of the *dsx* gene. The wild-type sequence is shown as reference. Amino acids that are different from the wild-type reference are highlighted in red.

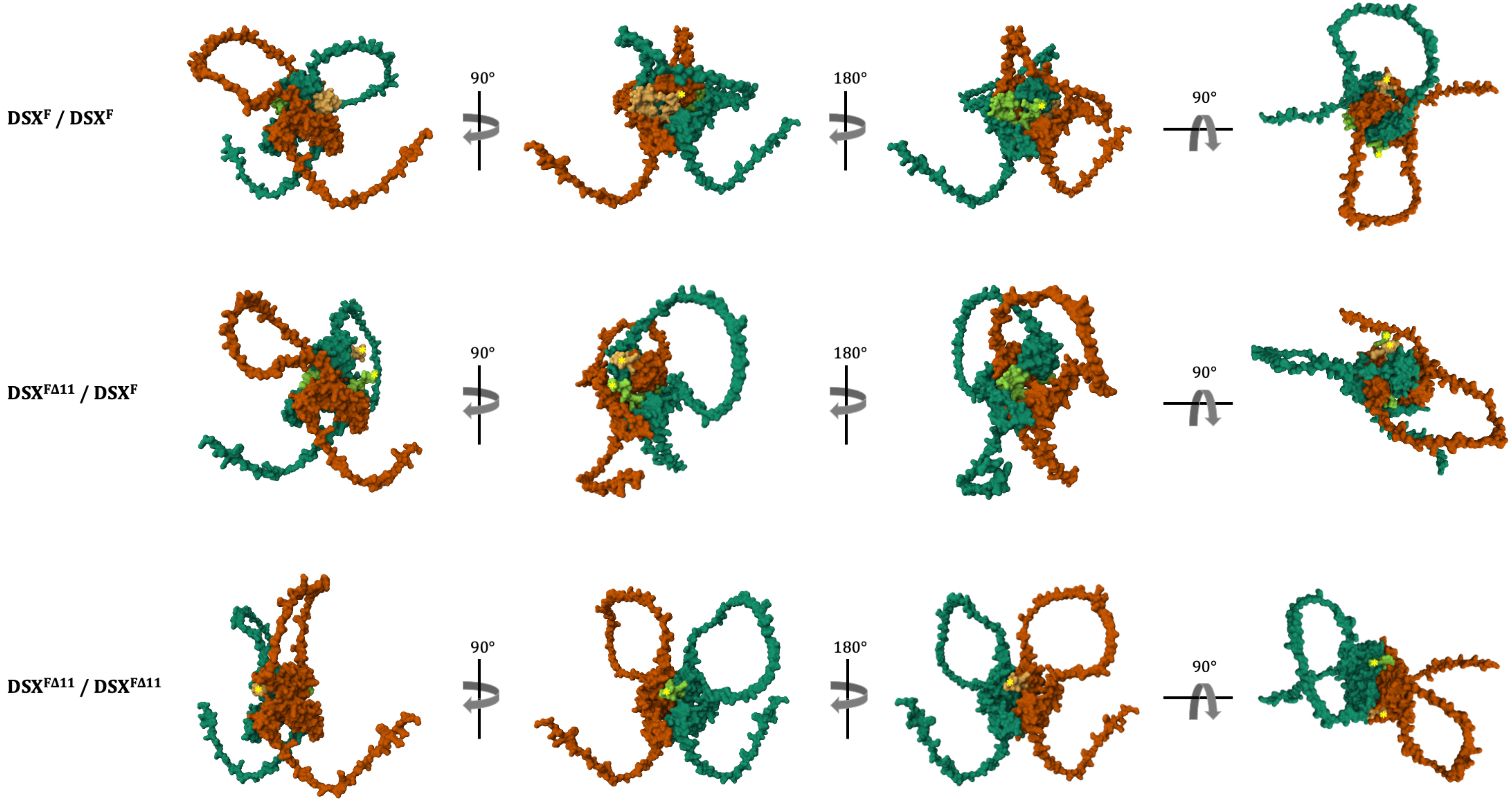


**Supplementary figure 3 – Predicted structure by Alpha-fold of the homodimers and heterodimers formed by DSX$^{F}$ and DSX$^{F\Delta11}$ proteins.** The top row shows the predicted structure of the wild-type DSX$^{F}$ homodimer (DSX$^{F}$/DSX$^{F}$), the middle one illustrates the predicted structure of the heterodimer (DSX$^{F}$/ DSX$^{F\Delta11}$) and the bottom row shows the predicted structure of the homodimer formed by two DSX$^{F\Delta11}$ proteins (DSX$^{F\Delta11}$/ DSX$^{F\Delta11}$). Monomers are represented in different colours, one in green and one in orange. Lighter colours in the structure illustrate the female specific region of DSX proteins, where the modifications by the *dsxF$^{\Delta11}$* allele are introduced. The C-terminal ends are indicated with a yellow star when shown from each perspective. In the heterodimer, the DSX$^{F}$ protein is represented in green, and the DSX$^{F\Delta11}$ one is shown in orange. While the OD1 domain (half bottom part in the first three columns) is generally conserved among the three dimers, considerable structural changes can be observed in the OD2 and especially in the female specific region of the proteins.

**Supplementary Table 1 – Frequencies of new mutations resulting from end joining repair mechanisms at the *dsx* target in the progenies of YLE$^{dsx}$; *dsxF*$^{GFP\text{-}null/+}$ males.** Each row corresponds to the progeny of a different mating couple (A-1 to A-8 are batches of progenies from YLE$^{dsx}$-A; *dsxF*$^{GFP\text{-}null/+}$ males, and B-1 to B-6 are batches of progenies from YLE$^{dsx}$-B; *dsxF*$^{GFP\text{-}null/+}$ males). The second column indicates the inheritance bias of the *dsxF*$^{GFP\text{-}null}$ allele (i.e, percentage of the offspring that inherited this allele) as a reference of the homing rates observed in each of these progenies. The third column shows the frequency of all newly generated mutations through end joining repair mechanisms (EJ) in the progeny that were absent in the progenitor male. The fourth column indicates the frequency of the most common mutation found in each progeny.

| Progeny | Inheritance of *dsxF*$^{GFP\text{-}null}$ allele (%) | Frequency of new EJ mutations (%) | Frequency of most common mutation (%) |
|---|---|---|---|
| **A-1** | 73.33 | 0.09 | 0.03 |
| **A-2** | 68.50 | 0.16 | 0.04 |
| **A-3** | 64.10 | 0.38 | 0.30 |
| **A-4** | 59.46 | 1.10 | 1.04 |
| **A-5** | 52.38 | 0.15 | 0.06 |
| **A-6** | 52.38 | 0.27 | 0.11 |
| **A-7** | 51.51 | 0.09 | 0.07 |
| **A-8** | 41.46 | 0.03 | 0.01 |
| **Average A** | - | **0.29** | **0.21** |
| **B-1** | 89.47 | 2.89 | 1.97 |
| **B-2** | 89.39 | 0.53 | 0.06 |
| **B-3** | 83.93 | 0.58 | 0.08 |
| **B-4** | 77.22 | 0.20 | 0.02 |
| **B-5** | 50.51 | 0.04 | 0.01 |
| **B-6** | 48.94 | 0.01 | 0.01 |
| **Average B** | - | **0.71** | **0.36** |

**Supplementary Table 2 – Frequencies of new mutations resulting from end joining repair mechanisms at the *dsx* target in the progenies of YLE$^{dsx}$-A; *dsxF*$^{\Delta11/+}$ males.** Amplicon sequencing was performed in pooled individuals of the progenies of YLE$^{dsx}$-A; *dsxF*$^{\Delta11/+}$ males. Each row corresponds to a different set of offspring. The second column indicates the frequency of the *dsxF*$^{\Delta11}$ allele in the progeny. The third column shows the frequency of the combination of all newly generated mutations through end joining repair mechanisms (EJ) found in each progeny. The fourth column indicates the frequency of the most common mutation (besides *dsxF*$^{\Delta11}$) in each progeny. Because a wild-type allele will always be inherited from the female, the percentage of individuals bearing each mutation corresponds to the double of the frequency of said mutation (i.e., if the frequency of the *dsxF*$^{\Delta11}$ allele is 50%, this reflects that 100% of the individuals bear the *dsxF*$^{\Delta11}$ allele). The sum of all mutations in progeny A-1 is >50%; this might be explained by either contamination or very low loads of nuclease deposition. Only in 3 progenies (A-1, A-2, A-4) there was a new mutation found in a frequency high enough to represent heterozygous individuals (0.59 – 2.14%).

| Progeny | Frequency of *dsxF*$^{\Delta11}$ allele (%) | Frequency of new EJ mutations (%) | Frequency of most common mutation (%) |
|---|---|---|---|
| **A-1** | 49.73 | 1.58 | 1.20 |
| **A-2** | 46.51 | 0.72 | 0.59 |
| **A-3** | 46.01 | 0.27 | 0.04 |
| **A-4** | 45.79 | 2.81 | 2.14 |
| **A-5** | 44.64 | 0.34 | 0.04 |
| **A-6** | 43.23 | 0.44 | 0.07 |
| **A-7** | 40.68 | 0.39 | 0.05 |
| **A-8** | 37.83 | 0.17 | 0.02 |
| **A-9** | 37.33 | 0.29 | 0.03 |
| **A-10** | 36.31 | 0.20 | 0.03 |
| **A-11** | 35.74 | 0.22 | 0.03 |
| **A-12** | 26.75 | 0.11 | 0.03 |
| **A-13** | 25.64 | 0.17 | 0.02 |
| **A-14** | 24.10 | 0.16 | 0.02 |
| **A-15** | 23.39 | 0.13 | 0.02 |
| **A-16** | 17.49 | 0.09 | 0.01 |
| **Average** | - | 0.51 | 0.27 |

**Supplementary Table 3 – Frequencies of new mutations resulting from end joining repair mechanisms at the *dsx* target in the progenies of YLE$^{dsx}$-B; *dsxF*$^{\Delta 11/+}$ males.** Amplicon sequencing was performed in pooled females of the progenies of YLE$^{dsx}$-B; *dsxF*$^{\Delta 11/+}$ males, separating the ones showing an intersex phenotype from those that had a wild-type phenotype. Each row corresponds to a different set of offspring. The second column shows the frequency of the *dsxF*$^{\Delta 11}$ allele in the intersex pool, and the third column indicates the frequency of all newly generated mutations through end joining repair mechanisms (EJ) in the male germline and inherited by females showing an intersex phenotype. The fourth column shows the frequency of the wild-type allele in the pool of females with a wild-type phenotype (expected to be 100% unless they inherited a newly generated recessive mutation), and the fifth column indicates the frequency of the aggregate of mutations in this pool. Because a wild-type allele will always be inherited from the female, the percentage of individuals bearing each mutation corresponds to the double of the frequency of said mutation (i.e., if the frequency of the *dsxF*$^{\Delta 11}$ allele is 50%, this reflects that 100% of the individuals bear the *dsxF*$^{\Delta 11}$ allele). Only one progeny (B-5) had a different mutation to *dsxF*$^{\Delta 11}$ in a high enough frequency to represent heterozygous individuals, found in the intersex pool, suggesting that it was a dominant negative mutation. Only one progeny (B-6) had a mutation in high frequency in the pool of wild-type-looking females. They were 2 individuals that were heterozygous for the *dsxF*$^{\Delta 11}$ allele, suggesting that they had inherited the allele from their progenitor and were either misclassified as wild-type or showed no intersex phenotype at pupal stage. (*) The average of mutations found in the wild-type pool does not include the progeny where the *dsxF*$^{\Delta 11}$ allele was found (more likely homed than created by MMEJ).

| Progeny | Freq of *dsxF*$^{\Delta 11}$ allele (%) – intersex pool | Freq of new EJ mutations (%) – intersex pool | Freq of wild-type allele (%) – wild-type pool | Freq of new EJ mutations (%) – wild-type pool |
|---|---|---|---|---|
| **B-1** | 50.05 | 0.05 | - | - |
| **B-2** | 50.87 | 0.07 | - | - |
| **B-3** | 51.49 | 0.07 | - | - |
| **B-4** | 50.23 | 0.12 | - | - |
| **B-5** | 37.51 | 14.26 | - | - |
| **B-6** | 50.88 | 0.06 | 49.07 | 50.93 |
| **B-7** | 46.51 | 0.07 | 99.99 | 0.01 |
| **B-8** | 51.33 | 0.05 | 99.99 | 0.01 |
| **B-9** | 51.39 | 0.06 | 99.98 | 0.02 |
| **Average** | - | 1.65 | - | 0.01* |

**Supplementary Table 4 – Model inheritance parameters and values for** YLE$^{dsx}$**.** Parameters that determine the inheritance of the released dominant mutation, the generation of new end-joining mutations (NHEJ) and sex-ratio distortion. The description of each parameter, the data source, and the values for the modelling of the YLE$^{dsx}$ are displayed.

| Parameter | Description | Data source | YLE$^{dsx}$ values |
|---|---|---|---|
| **Gamete production** | | | |
| $\boldsymbol{d}$ | Proportion of offspring which carry the desired mutation | Experiment | 0.945 |
| $\boldsymbol{v}$ | Proportion of offspring which carry an alternative mutation | Experiment | 0.020 |
| $\boldsymbol{e}$ | Probability of homing | $2d-1$ | 0.890 |
| $\boldsymbol{u}$ | Proportion of non-homed chromosomes which are NHEJ | $\frac{\mathrm{v}}{1-d}$ | 0.455 |
| $\boldsymbol{c}$ | Probability of cleavage (in heterozygotes) | $e+(1-e)u$ | 0.930 |
| $\boldsymbol{j}$ | Probability of NHEJ given joining (in heterozygotes) | $\frac{(1-e)u}{e+(1-e)u}$ | 0.043 |
| $\boldsymbol{\mu}$ | Probability of mutation (in wild-type homozygotes) | Unknown | 1 |
| $\boldsymbol{p}$ | Proportion of dominant NHEJ (in heterozygotes) | Unknown | 0 |
| $\boldsymbol{q}$ | Proportion of dominant NHEJ (in wild-type homozygotes) | Unknown | 0 |
| $\boldsymbol{m}$* | Proportion of Y-bearing sperm produced by males carrying an X-shredder | Pollegioni et al 2020 | 0.9 |

*Relevant only when modelling the YLE in the presence of an X-shredder.

**Supplementary Table 5 – Model fitness parameters.** Parameters that determined the fitness effects of the autosomal alleles $\boldsymbol{\alpha}$ and $\alpha$ relative to the WT allele A, and of the X-shredder. The description of each parameter and the values used for each strategy modelled is displayed.

| **Fitness costs** | | **YLE$^{dsx}$** | **YLE$^{dsx}$ + XS** | **SIT and RIDL** | **fsRIDL** | **XS** |
|---|---|---|---|---|---|---|
| $\boldsymbol{s_{\alpha}^{G}}$ | Fitness cost for the dominant cleavage resistant allele | $1^F$ | $1^F$ | 1 | $1^F$ | - |
| $\boldsymbol{s_{a}^{G}}$ | Fitness cost for the recessive cleavage resistant allele | $1^F$ | $1^F$ | - | - | - |
| $\boldsymbol{h_{\alpha}^{G}}$ | Dominance coefficient for the dominant cleavage resistant allele | $1^F$ | $1^F$ | 1 | $1^F$ | - |
| $\boldsymbol{h_{a}^{G}}$ | Dominance coefficient for the recessive cleavage resistant allele | 0 | 0 | - | - | - |
| $\boldsymbol{s_{a\alpha}^{G}}$ | Fitness cost for heterozygous for the two types of cleavage resistant allele | $1^F$ | $1^F$ | - | - | - |
| $\boldsymbol{s_{XS}^{M}}$ | Fitness cost for the X-shredder allele in males | - | 0.2* | - | - | 0 |

The superscript $^F$ indicates the case where fitness costs are applied only to females and the value in males is zero.
*Pollegioni et al 2020.
